# Factor XI localization in human deep venous thrombus and function of activated factor XI on venous thrombus formation and hemostasis in rabbit

**DOI:** 10.1101/2023.09.19.558346

**Authors:** Nobuyuki Oguri, Toshihiro Gi, Eriko Nakamura, Kazunari Maekawa, Eiji Furukoji, Hoshimi Okawa, Sho Kouyama, Saki Horiuchi, Tatefumi Sakae, Minako Azuma, Yujiro Asada, Atsushi Yamashita

**Affiliations:** Department of Pathology, Faculty of medicine, University of Miyazaki, Miyazaki, Japan; Department of Radiology, Faculty of medicine, University of Miyazaki, Miyazaki, Japan; ONO Pharmaceutical Co., Ltd., Osaka, Japan; Department of Diagnostic Pathology, Miyazaki Medical Association Hospital, Miyazaki, Japan

**Keywords:** coagulation factor x, coagulation factor xi, hemostasis, pathology, venous thromboembolism

## Abstract

**Background:** Novel anticoagulants targeting coagulation factor XI (FXI)/activated FXI (FXIa) are under development. However, whether FXI is present in human deep vein thrombosis (DVT) and whether FXIa and activated factor X (FXa) play different roles in venous thrombus formation and hemostasis remain unclear. This study aimed to determine the presence of FXI in DVT and the effects of direct oral FXIa and FXa inhibitors on venous thrombus formation and hemostasis in rabbits and mural thrombus formation in flow chamber system.

**Methods:** We immunohistochemically assessed FXI localization in human aspirated DVT (n=15). Additionally, we compared thrombus formation induced by endothelial denudation and stenosis in jugular vein, and skin bleeding time and volume between rabbits treated with direct FXIa inhibitors (ONO-1600586) and FXa inhibitors (rivaroxaban). *Ex vivo* rabbit and human blood were perfused on a flow chamber under low shear rates (70/s).

**Result:** FXI was localized in all DVT, predominantly in fibrin-rich areas. The FXI-immunopositive area in the non-organizing area was greater than that in the organizing area. Although FXIa and FXa inhibitors comparably inhibited venous thrombus formation, FXIa inhibitors did not affect bleeding time or volume in rabbits. FXIa or FXa inhibitors mildly or strongly inhibited fibrin formation at low shear rates respectively. Furthermore, the FXIa inhibitor suppressed human FXIa activity, thrombin generation, and fibrin formation during perfusion.

**Conclusion:** The pathological findings of human DVT suggest FXI’s role in human DVT. FXIa inhibitors may inhibit less fibrin formation than FXa inhibitors, and may explain the minor role of FXIa in hemostasis.

**Essential:**

- Presence of factor XI (FXI) in venous thrombus and less bleeding in its inhibition are unclear.
- We assessed FXI localization in deep vein thrombosis (DVT) and function of FXIa in rabbit.
- FXI localized in human DVT that provide a rationale for FXI inhibition in human DVT.
- FXIa inhibitor inhibited less fibrin formation than factor Xa inhibitor under low-shear rate.

## Introduction

Deep vein thrombosis (DVT) and pulmonary embolism (PE), which are collectively termed venous thromboembolism (VTE), constitute major global health challenges. The estimates of VTE incidence ranged from 79 to 269 per 100,000 population [1]. Current oral anticoagulants that inhibit the protease thrombin or activated factor X (FXa) or lower the plasma concentrations of their precursors, prothrombin or factor X (FX), can reduce the risks of VTE and ischemic stroke. However, their use has a certain risk of major or minor bleeding [2]. Even if the patients with atrial fibrillation receive low-dose direct oral anticoagulant, the incidence of major bleeding accounts for approximately 3% per year [3]. Therefore, developing anticoagulants without or with fewer bleeding complications is crucial.

Factor XI (FXI) is a serine protease in the intrinsic pathway of coagulation, which is activated by thrombin, activated FXI (FXIa), and activated factor XII (FXIIa). Elevated plasma FXI levels are associated with VTE and ischemic stroke [4,5]. Furthermore, people with congenital FXI deficiency have a lower risk for cardiovascular and VTE events [6,7]. The age-adjusted incidence rate of VTE was two-thirds lower in individuals with FXI deficiency (FXIa activity ≤ 50 %) than in those with normal FXIa activity [7]. In animal models, a genetic deficiency of FXI or inhibition of FXI and/or FXIa by antisense oligonucleotide, antibody, and small molecules reduced venous thrombus weight without increasing the bleeding time [8–11]. However, ferric chloride [8,10] and silk thread [9] are not physiological initiators of venous thrombus formation, and a small molecule FXIa inhibitor was intravenously administrated [10]. Therefore, it has not been reported whether there are any differences in venous thrombus formation and bleeding between oral administration of a small molecule FXIa and FXIa inhibitors. In addition, the exact mechanisms of the less bleeding phenotype in FXI and FXIa inhibition are not clearly understood. Furthermore, the presence of FXI in human DVT has not yet been reported.

Therefore, this study aimed to investigate FXI localization in human DVT, the function of FXIa and FXa in venous thrombus formation and hemostasis in rabbits, and mural thrombus formation in a flow chamber system.

## Methods

### Aspirated thrombi from patients with DVT

The Ethics Committee of the University of Miyazaki approved the study protocol (approval no. O-0684). Overall, 15 thrombi were obtained from 15 patients (8 men and 7 women; age range, 20–78 years; median, 54 years; Table S1) with DVT who were diagnosed based on clinical symptoms, laboratory data, and clinical imaging findings. Thrombi were aspirated from the proximal portion using a guiding catheter (Guider Softip Guiding Catheter; Boston Scientific Japan, Tokyo, Japan). This catheter was placed from the popliteal vein to the iliac vein (10 cases) and the leg vein to the inferior vena cava (5 cases). X-ray venography revealed an extensive filling defect in the veins before thrombus aspiration and a reduced filling defect after thrombus aspiration. Finally, all aspirated thrombi were immediately fixed in 4% paraformaldehyde and embedded in paraffin for histological evaluation.

### Histological findings of aspirated DVT

Briefly, 4-μm thick sections were stained with hematoxylin and eosin (HE) and morphologically assessed. Because the venous thrombi showed time-dependent changes, including fresh components, cell lytic changes, and organizing reactions. We defined the fresh component and cell lytic change as non-organizing area or the proliferation of endothelium-like flat cells and fibroblastic cells and fibrous matrix deposition as organizing area [12,13]. Endothelialization was confirmed through immunohistochemistry for CD34 [14]. Finally, we compared the immunopositive areas described below between the fresh area and organizing area in each thrombus.

### Immunofluorescence of venous thrombus

Immunofluorescence was performed to examine FXI localization in DVT. The DVT sections were stained with an anti-FXI antibody (sheep polyclonal, LS-B10243; LifeSpan BioSciences (LSBio), Inc. Seattle, USA), anti-fibrin antibody (mouse monoclonal, clone 59D8; EMD Millipore Corp., Temecula, CA, USA), anti-von Willebrand factor (VWF) antibody (rabbit polyclonal, Dako/Agilent, Santa Clara, CA, USA), or anti-glycophorin A antibody (mouse monoclonal, clone JC159; Dako/Agilent). CF488 conjugated-donkey anti-rabbit IgG (Biotium, Hayward, CA, USA) and CF568 conjugated-donkey anti-mouse IgG (Biotium, Hayward) were used as secondary antibodies. Figure S1 shows an immunopositive band for human FXI (HFXI 1111, Enzyme Research Laboratories Ltd., Swansea, UK) but not human prekallikrein (HPK 1302, Enzyme Research Laboratories Ltd.) in Western blotting using the anti-FXI antibody (LS-B10243; LSBio).

### Immunohistochemistry for human DVT

Briefly, consecutive 4-μm slices of human aspirated thrombi were immunohistochemically stained using antibodies against CD34 (mouse monoclonal, clone QBEnd10; Dako/Agilent), FXI (LSBio), fibrin (Millipore), anti-platelet glycoprotein (GP) IIb/IIIa (Sheep monoclonal, Affinity Biologicals Inc., Ancaster, Canada), glycophorin A (Dako/Agilent), and CD66b (mouse monoclonal, clone 6/40c; BioLegend, San Diego, CA, USA). Next, the sections were stained with EnVision anti-mouse, rabbit immunoglobulin (Dako/Agilent), or anti-sheep secondary antibodies (Jackson ImmunoResearch Inc. West Grove, PA, USA). Horseradish peroxidase activity was visualized using 3, 3’-diaminobenzidine tetrahydrochloride, and the sections were mildly counterstained using Meyer’s hematoxylin. Immunostaining controls included non-immune mouse IgG or non-immune sheep serum rather than primary antibodies. Microscopic images were captured using a photosensitive color CCD camera (DS-Fi3, Nikon, Tokyo, Japan) under a 40 × objective lens. The immunopositive areas for FXI, fibrin, glycophorin A, and GPIIb/IIIa in the non-organizing and organizing areas were semi-quantified using a color imaging morphometry system (WinROOF, Mitani, Fukui, Japan). The detailed method has been described in a previous study [13]. Briefly, immunopositive areas were extracted as green areas using specific protocols based on the color parameters of hue, lightness, and saturation. The data were expressed as the mean ratio of positively stained areas per thrombus area in three fields [13]. Furthermore, the number of CD66b immunopositive cells was counted in the five most densely stained fields under a 40× objective lens, and cell density was expressed as the number of immunopositive cells per mm^2^.

### Oral direct activated FXI inhibitor and *in vitro* enzyme assay

We used the FXIa inhibitor ONO-1600586 (ONO Pharmaceutical Co. Ltd., Osaka, Japan), which is a low molecular weight compound (a molecular weight of 507), to investigate the role of FXIa in venous thrombus formation and hemostasis. The inhibitory activity of the compounds against FXIa, FXa, FXIIa, activated factor IXa (FIXa), activated factor VII (FVIIa), plasma kallikrein, and thrombin was evaluated using the appropriate purified proteases and synthetic substrates. Furthermore, the hydrolysis rate of the chromogenic substrate by relevant proteases was continuously measured at 405 nm. Moreover, the inhibitory activity against each enzyme was calculated as % inhibition using the equation described below: % Inhibition = [[(rate without compound)-(rate with compound)]/(rate without compound)] ×100%.

Each half-maximal inhibitory concentration (IC_50_) value was determined by plotting the concentration of the compound against the % inhibition.

Human FXIa (Haematologic Technologies Inc. Essex Junction, VT, USA) activity was measured at an enzyme concentration of 0.1 U/mL in 150 mM NaCl (FUJIFILM Wako Pure Chemical Corp., Osaka, Japan), 5 mM KCl (FUJIFILM Wako Pure Chemical Corp.), 1 mg/mL polyethylene glycol (PEG6000, FUJIFILM Wako Pure Chemical Corp.), and 50 mM HEPES (Dojindo Laboratories, Kumamoto, Japan)-NaOH (Nacalai Tesque Inc., Kyoto, Japan) (pH 7.4) with 300 μM L-pyroglutamyl-L-prolyl-L-argininep-nitroaniline hydrochloride (Chromogenix S-2366, Diapharma, West Chester, OH, USA).

Human plasma kallikrein (Enzyme Research Laboratories Ltd.) activity was measured at an enzyme concentration of 0.605 mU/mL in 200 mM NaCl, 5 mg/mL PEG6000, and 100 mM Phosphate (FUJIFILM Wako Pure Chemical Corp.)-NaOH (pH 7.4) with 150 μM H-D-prolyl-L-phenylalanyl-Larginine-p-nitroaniline dihydrochloride (Chromogenix S-2302, Diapharma).

Human FXa (American Diagnostica Inc. Stamford, CT, USA) and thrombin (Sigma-Aldrich Co., LLC., St. Louis, MO, USA) activities were measured at the enzyme concentrations of 0.18 U/mL and 0.12 U/mL, respectively in the same buffer containing 150 mM NaCl, 2 mg/mL PEG6000, and 50 mM Tris (Nacalai Tesque Inc.)-HCl (Nacalai Tesque Inc.) (pH 7.4), except that the reactions were started with 300 μM N-benzoyl-L-isoleucyl-L-glutamyl-glycyl-Larginine-p-nitroaniline hydrochloride and its methyl ester (Chromogenix S-2222, Diapharma) and 300 μM Chromogenix S-2366 (Diapharma).

Human factor α-XIIa (Enzyme Research Laboratories Ltd) activity was measured at an enzyme concentration of 0.17 U/mL in 150 mM NaCl and 50 mM Tris-HCl (pH 7.4) with 300 μM Chromogenix S-2302 (Diapharma).

Human FIXa (American Diagnostica Inc.) activity was measured at an enzyme concentration of 13 U/mL in 100 mM NaCl, 5 mM CaCl_2_ (Nacalai Tesque Inc.), 30% ethylene glycol (Kanto Chemical Co., Inc., Tokyo, Japan), and 50 mM Tris-HCl (pH 7.4) with 3 mM Pefachrome FIXa 3960 (Leu-Ph’Gly-Arg-pNA, Pentapharm Ltd., Basel-Landschaft, Switzerland).

Human FVIIa activity was measured using recombinant human FVIIa (American Diagnostica Inc.) in the presence of recombinant human tissue factor, which was produced at ONO Pharmaceutical Co. Ltd., Osaka, Japan, in a buffer containing 150 mM NaCl, 5 mM CaCl_2_, 0.5 mg/mL PEG6000, and 50 mM HEPES-NaCl (pH 7.4) with 3 mM H-D-isoleucyl-L-prolyl-L-arginine-p-nitroaniline dihydrochloride (Chromogenix S-2288, Diapharma).

### Rabbit model of jugular vein thrombosis and skin bleeding

The Animal Care Committee of the University of Miyazaki approved the animal study protocols (Approval No. 2019-532), which conformed to the Guide for the Care and Use of Laboratory Animals published by the U.S. National Institutes of Health. Overall, 71 healthy male Japanese white rabbits, purchased from Japan SLC Incorporation (Shizuoka, Japan), weighing 2.8–3.3 kg, were fed with a conventional diet.

The protocols of the venous thrombosis and skin bleeding models in rabbits were as follows (Figure S2). First, fasting rabbits were orally administered a solvent as a control (n=10), an FXIa inhibitor (ONO-1600586, 5 mg/kg n=10, 15 mg/kg n=10, and 50 mg/kg n=11), or an FXa inhibitor (rivaroxaban, 1.5 mg/kg, 5 mg/kg, and 15 mg/kg, n=10 each) with a Nelaton catheter of 5mm diameter (IZUMO Health Co. Ltd., Nagano, Japan) 90 min before thrombus formation. The thrombus formation and bleeding were performed under general anesthesia through the subcutaneous injection of a mixture of medetomidine hydrochloride (0.08 mg/kg body weight), butorphanol tartrate (0.2 mg/kg), and midazolam (0.4 mg/kg) [15]. Additional anesthesia with the one-third amount of the above-mentioned mixture of drugs was injected 80 min after the first injection, and all surgical operations proceeded under aseptic conditions.

### Thrombus formation in rabbit jugular veins by endothelial denudation and luminal stenosis

Briefly, 90 min post-administration of the solvent, ONO-1600586, or rivaroxaban, venous thrombi were induced in rabbit jugular veins through endothelial denudation using a 3F balloon catheter (Edwards Lifesciences, Irvine, CA, USA) according to a previous study [15], and the method was modified to induce luminal stenosis. A 2-mm polyethylene tube was placed outside the vessel. Both the jugular vein and polyethylene tube were ligated, and the tube was removed (Figure S3) [16]. Next, 3 hours after thrombus formation and stenosis, the rabbits were infused with heparin (500 U/kg, i.v.) and then sacrificed with an overdose of pentobarbital (60 mg/kg, i.v.). Subsequently, the animals were perfused with 50 mL phosphate-buffered saline (0.01 mol/L), and the jugular veins were sampled. The venous thrombi were immediately removed from the venous wall, and weighed. Finally, the thrombus was fixed in 4% paraformaldehyde for 24 hours and embedded in paraffin for histological evaluation.

### Bleeding time and volume by skin incision in rabbit

Ninety minutes after administering the solvent, rivaroxaban (1.5, 5, and 15 mg/kg), and ONO-1600586 (5, 15, and 50 mg/kg), skin bleeding was induced using two lines of 3 cm length incisions on both legs of the rabbit. Next, wet Kimwipes (Kimberly-Clark Corp., TX, USA) were placed on the injured lesions, and the bleeding time was measured. To measure the bleeding volume, we collected the Kimwipes into a falcon tube filled with 20 mL of saline each 1–2 min for 30 min and then removed the Kimwipes. The collected blood was centrifuged at 900 × g for 5 min at room temperature, and the supernatant was discarded. One mL of distilled water was added to the pellet and then hemolyzed. The bleeding volume was estimated using a spectrophotometer at 540 nm (SpectraMax 190 Microplate Reader, Molecular Devices, LLC., San Jose, CA, USA). A standard curve was generated through the sequential dilution of rabbit blood. Finally, the number of erythrocytes was measured using a hematology analyzer (XP-300, Sysmex, Kobe, Japan).

### Histology and immunohistochemistry of jugular vein thrombus

Paraffin-embedded sections (4-µm thick) of the rabbit jugular vein thrombus were stained with HE and morphologically assessed. Consecutive sections of thrombi were immunohistochemically stained with antibodies against fibrin (Millipore) and GPIIb/IIIa (Affinity Biologicals Inc.). Images were digitized using a photosensitive color CCD camera (BX51, OLYMPUS). Finally, the area of the red-stained erythrocytes in HE staining and the immunopositive areas for fibrin and GPIIb/IIIa were semi-quantified using a color imaging morphometry system (WinROOF, Mitani).

### Coagulation parameters

Blood samples were collected from the central ear arteries of rabbits before, 75 min, and 4.5 hours after the administration of the solvent, ONO-1600586 and rivaroxaban, in 3.8% sodium citrate (9:1 v/v) using 21-G needles. Plasma samples were centrifuged at 1,500 g for 15 min at room temperature. Plasma prothrombin time (PT) and activated partial thromboplastin time (aPTT) were measured using a coagulation timer (KC-1 Delta, Tcoag Ireland Ltd., Wicklow, Ireland) and reagents (aPTT: ACTIN, Siemens, Munich, Germany; PT: THROMBOCHECK PT+, Sysmex, Kobe, Japan). Data were expressed as the ratio of PT and aPTT to those before administration.

### *Ex vivo* rabbit blood perfusion with flow chamber system and visualization of the thrombi on the collagen surface

Blood from rabbits 75 min postadministration of solvent as a control (n=5), an FXIa inhibitor (ONO-1600586, 50 mg/kg n=5), or an FXa inhibitor (rivaroxaban, 15 mg/kg n=5) was collected using 21-G needles into plastic syringes containing corn trypsin inhibitor (Molecular Innovations, Southfield, MI; final concentration 20 µg/mL) and a specific FXa inhibitor, Arixtra, (Organon Sanofi-Synthelabo, West Orange, NJ; final concentration 6 µg/mL). Anticoagulated whole blood was stored at room temperature and used for perfusion studies within 1 hour of collection. Next, mepacrine (quinacrine dihydrochloride, Sigma-Aldrich; final concentration 5 µM) was added to the blood before perfusion for platelet labeling, which enables the visualization of platelet-surface interaction through epifluorescence videomicroscopy [11]. Collagen from horse tendon (Collagen Reagent HORM, Takeda Pharmaceutical Co., Ltd., Tokyo, Japan, 20 µg/mL) was immobilized on glass coverslips (24×50 mm) in parallel-plate flow chambers as described [11]. Blood samples were perfused through the chamber using a syringe pump (KDS100, KD Scientific Inc., New Hope, PA) at a constant flow rate (15 mL/hour) to achieve wall shear rates of 70/s. Platelets interacting with the immobilized collagen were visualized using an inverted-stage epifluorescence microscope (IX71, OLYMPUS, Tokyo, Japan). Images were captured at 3, 6, and 9 min after perfusion with a photosensitive color CCD camera (DP70, Olympus), and surface-covering areas were semi-quantified using a morphometric analysis system (WinROOF, Mitani).

After perfusion, coverslips were immediately fixed in 95% alcohol for 1 hour at room temperature, stained using antibodies for fibrin (Millipore), and the coverslips were faintly counterstained with Meyer’s hematoxylin. The microscopic images were digitized using a photosensitive color CCD camera (DS-Fi3, Nikon), and the immunopositive areas of the images were semi-quantified for each antibody using a color imaging morphometry system (WinROOF, Mitani). Finally, the number of attached cells with multilobulated nuclei was counted as the neutrophil number in five fields under a 20 × objective lens and expressed as the number of neutrophils per mm^2^.

### *In vitro* human blood perfusion and measurement of blood coagulation before and after the perfusion

The Ethics Committee of the University of Miyazaki approved this study (approval no. O-1160). The Collagen Reagent HORM (Takeda, 20 µg/mL) was immobilized on glass coverslips (24×50 mm) in parallel-plate flow chambers as described above. Blood from healthy human volunteers who were not on medication was collected using 21-G needles into plastic syringes containing a corn trypsin inhibitor (Molecular Innovations; final concentration 10 µg/mL). Anticoagulated whole blood was used for the perfusion. The solvent (n=5) or an FXIa inhibitor (ONO-1600586, 3 µmol/L n=5) and mepacrine (Sigma-Aldrich; final concentration 5 µM) were added to the blood before perfusion. Blood samples were perfused through the chamber using a syringe pump (KDS100) at a constant flow rate (15 mL/hour) to achieve wall shear rates of 70/s. Finally, the surface-covering area, fibrin-immunopositive area, and the number of neutrophils were analyzed as described above.

The pre– and post-perfused blood samples were collected into a 3.8% sodium citrate anticoagulant (9:1 v/v). Plasma samples were centrifuged at 1,500 g for 15 min at room temperature. FXIa and FXa activities were measured using an automatic blood coagulation analyzer (ACL-TOP, Werfen Headquarters, Barcelona, Spain) as well as FXI-deficient plasma (HemosIL Factor XI Deficient Plasma, Werfen, HemosIL Synthesil APTT, Werfen) and FX-deficient plasma (HemosIL Factor X Deficient Plasma, Werfen, HemosIL RecombiPlasTin, Werfen). The plasma levels of prothrombin fragment 1+2 were measured using an enzyme-linked immunosorbent assay (Enzygnost F1+2 monoclonal, Siemens).

### Statistical analyses

Data analysis was performed using GraphPad Prism 9 (GraphPad Software Inc., California, USA). All procedures were performed using a blinded experimental design. All data are expressed as means ± standard deviation (SD) or median and interquartile range when the variance was skewed. Differences between or among individual groups were statistically evaluated using the Mann–Whitney U-test, Kruskal–Wallis test with Dunn’s multiple comparison test, or a one-or two-way analysis of variance (ANOVA) with Tukey’s multiple comparison test as appropriate. Statistical significance was considered at P˂0.05 (n indicates the number of samples). We have indicated the statistics in the legends of each figure.

## Results

### Localization of FXI in human DVT

All 15 thrombi were immunopositive for FXI. Immunofluorescent images showed that FXI was closely localized to fibrin rather than VWF or glycophorin A (Figure 1A). We measured the immunopositive areas for FXI, fibrin, platelets, erythrocytes, and neutrophils to examine the extent of thrombus components and FXI in the non-organizing and organizing areas. The organizing area was confirmed by the presence of CD34 immunopositive cells, which correspond to endothelialization. All 15 thrombi were immunopositive for glycophorin A (an erythrocyte marker), GPIIb/IIIa (a platelet marker), fibrin, and CD66b (a neutrophil marker) (Figure 1B). The areas immunopositive for FXI, fibrin, platelets, erythrocytes, and CD66b in the non-organizing area were larger than those in the organizing area (Figure 1C).

**Figure 1.**
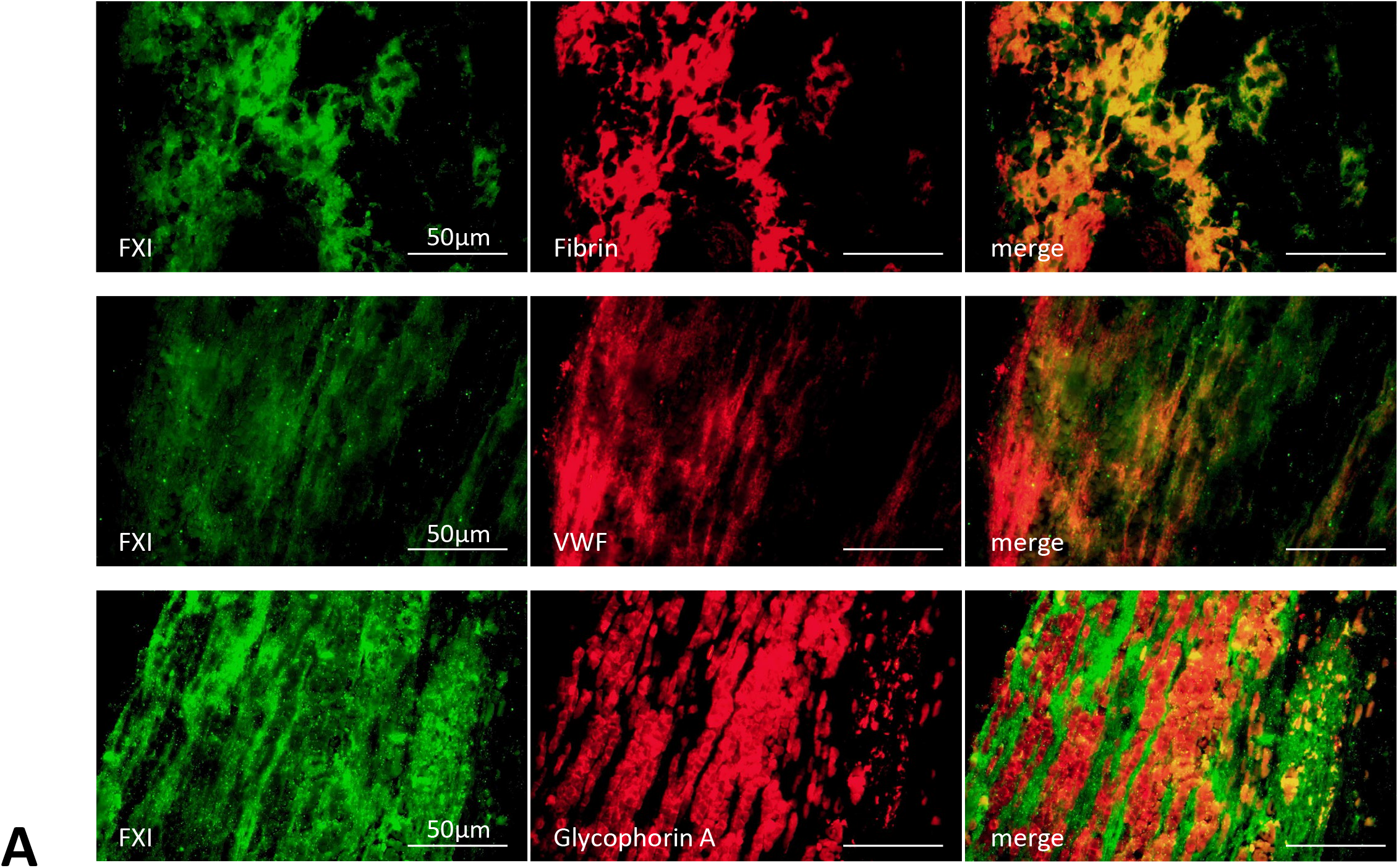
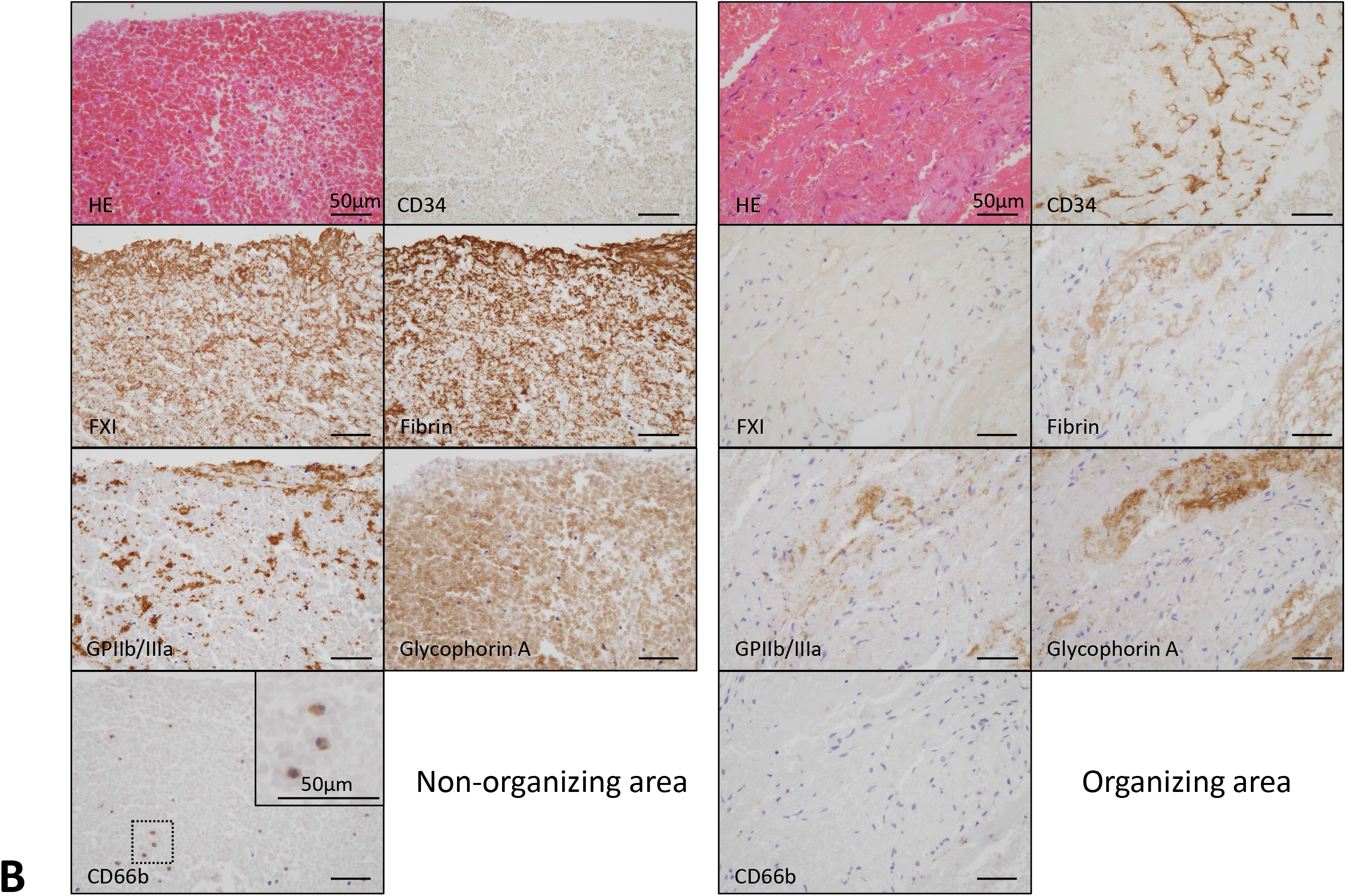
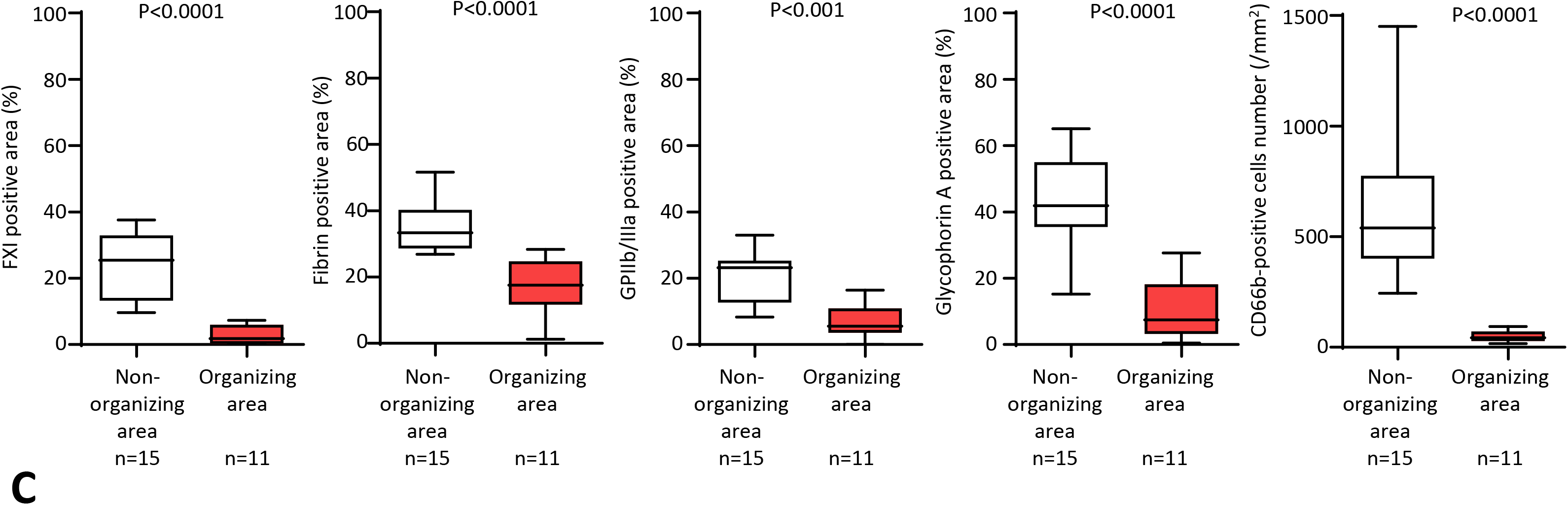
Presence of factor XI (FXI) in human deep vein thrombosis (DVT) **A**. Representative immunofluorescent images of fresh component of DVT. Upper row shows CF488-labeled FXI (green), CD568-labeled fibrin (red), and merged images. Middle row shows CF488-labeled FXI, CD568-labeled von Willebrand factor (VWF), and merged images. Lower row shows CF488-labeled FXI, CD568-labeled glycophorin A, and merged images. FXI was closely localized with fibrin, and partly localized with VWF, a marker of platelets. Glycophorin A, a marker of erythrocytes, is predominantly surrounded by FXI. **B**. Representative immunohistochemical images in fresh and organizing areas of DVT. The organizing areas were confirmed by presence of CD34 immunopositive cells, corresponding to endothelialization/ organization, while the fresh area was confirmed by absence of CD34 immunopositive cells. FXI localized fibrin– and erythrocyte-rich fresh area. In the fresh area, aggregated clusters of platelets and sparsely distributed neutrophils are present. Inset in CD66b represents a high-magnification image of the dashed square. In the organizing area, immunoreaction for FXI, fibrin, glycoprotein IIb/IIIa (GPIIb/IIIa), glycophorin A, or CD66b is modest or focal. **C**. Immunopositive area for FXI, fibrin, GPIIb/IIIa, and glycophorin A, and CD66b-immunopositive cell density in non-organizing (CD34-immunonegative) and organizing (CD34-immunopositive) areas. (Mann-Whitney U-test)

### Characterization of an activated factor XI inhibitor (ONO-1600586)

We used an activated FXI inhibitor, ONO-1600586, to evaluate the function of FXIa. ONO-1600586 prolonged aPTT but did not affect PT in both humans and rabbits. Additionally, ONO-1600586 concentration at aPTT doubling time was 0.60 µM and 0.61 µM for humans and rabbits, respectively. ONO-1600586 did not double PT at 33 µM in plasma concentration for humans and rabbits. ONO-1600586 inhibited human FXIa (IC50:0.0020 µM) and weakly human kallikrein (IC50:0.12 µM) but did not inhibit human thrombin, FVIIa, FIXa, FXa, FXIIa, plasmin, urokinase, or t-PA (IC50: >25 µM) (Table S2).

### Oral administration of FXIa inhibitor prolongs aPTT rather than PT in rabbits, whereas oral administration of FXa inhibitor prolongs both PT and aPTT

We evaluated PT and aPTT before and after the oral administration of the solvent, an FXIa inhibitor (ONO-1600586), and an FXa inhibitor (rivaroxaban) during venous thrombus formation and measured the bleeding time. ONO-1600586 (15 and 50 mg/mL) dose-dependently prolonged aPTT rather than PT to 75 min and 4.5 hours. Rivaroxaban (5 and 15 mg/mL) dose-dependently prolonged both PT and aPTT to 75 min and 4.5 hours. However, oral administration of the solvent did not affect the PT or aPTT at either time point (Figure 2A and B).

**Figure 2.**
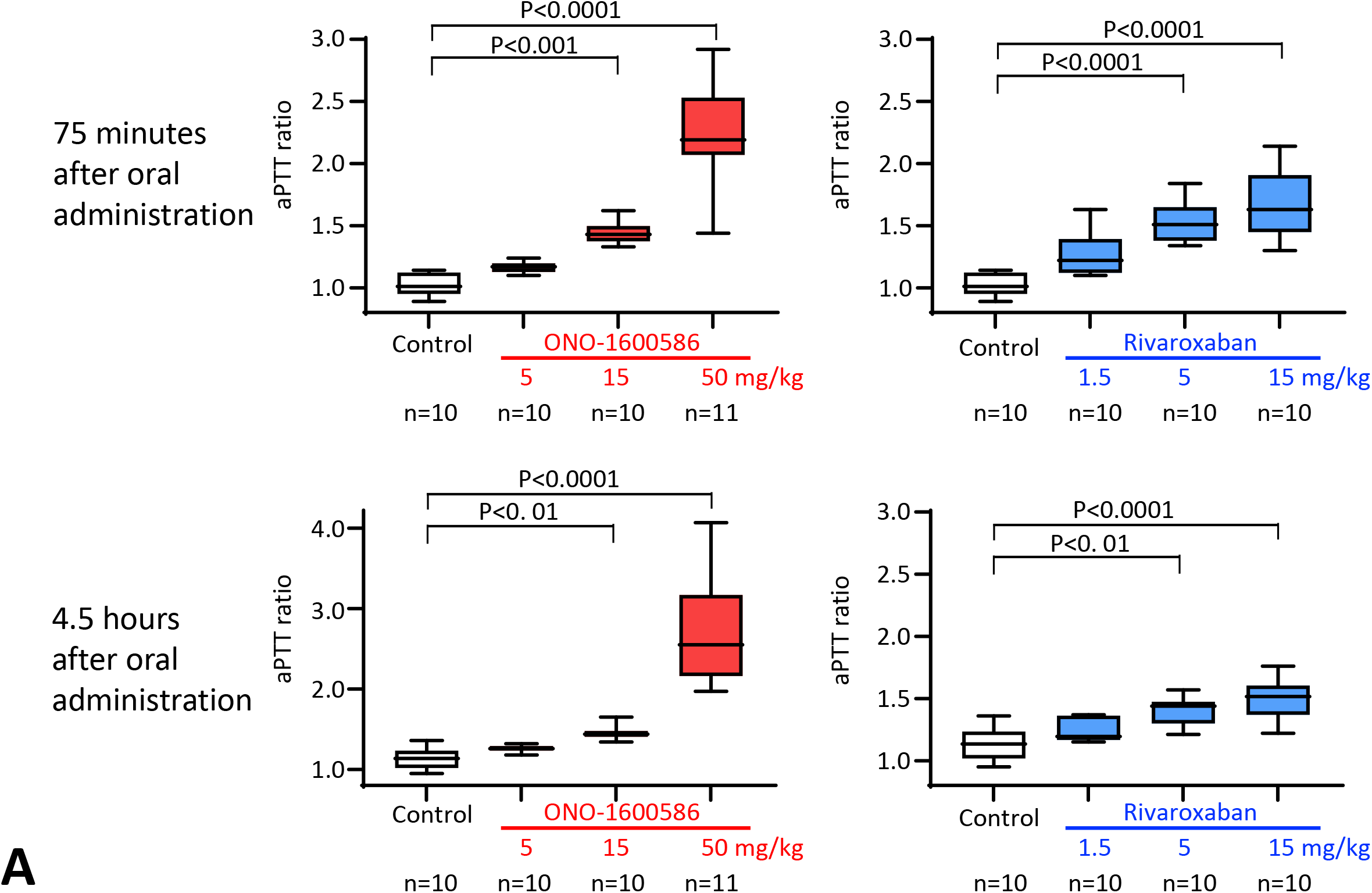
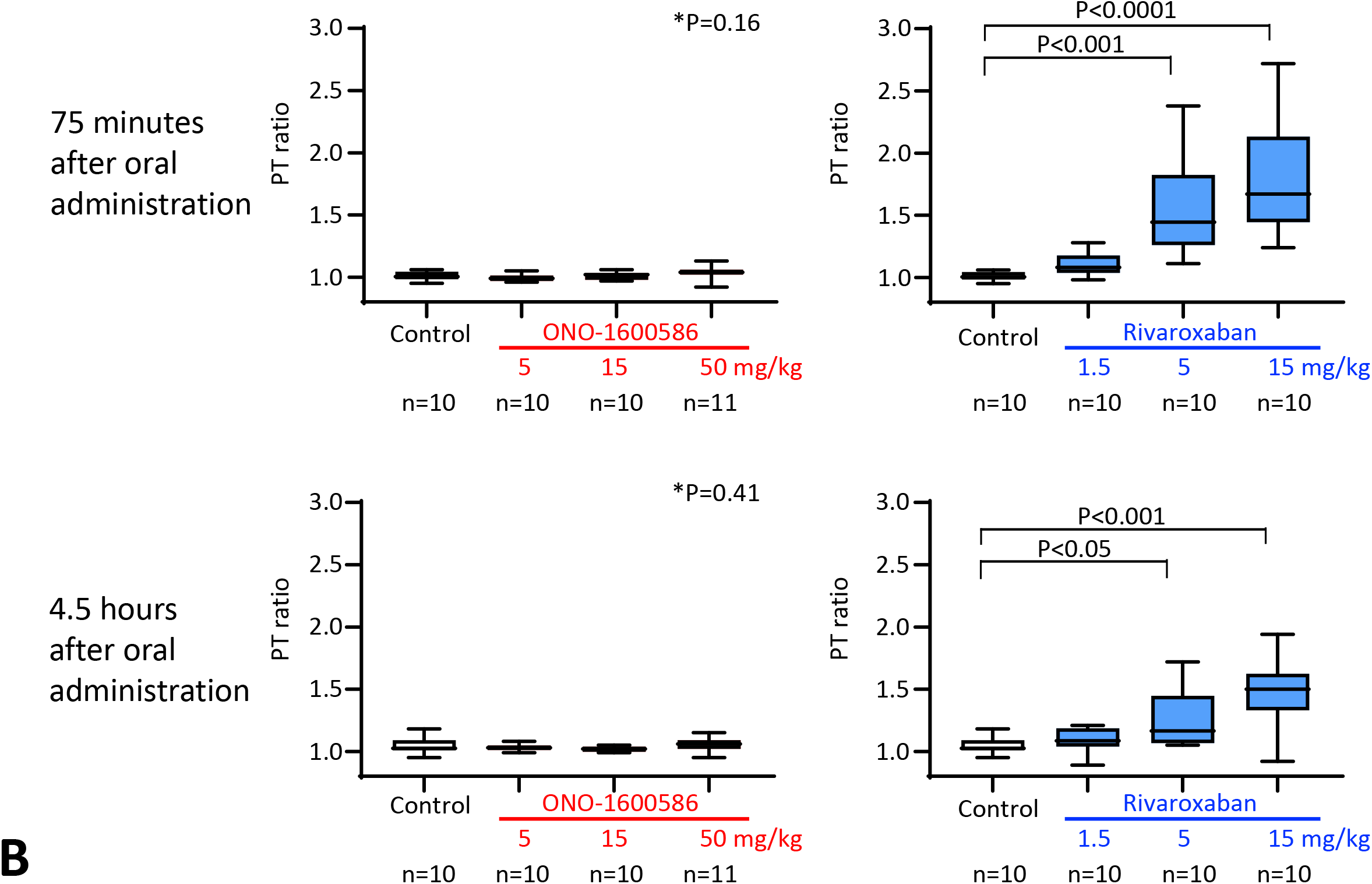
Activated partial thromboplastin time (aPTT) and plasma prothrombin time (PT) ratio after oral administration of ONO-1600586 and rivaroxaban. **A**. aPTT ratio 75 minutes (before thrombus formation and bleeding test) and 4.5 hours (3 hours after thrombus formation) after oral administration of solvent (control) and activated factor XI (FXIa) (ONO-1600586) and factor X (FX) inhibitors (Rivaroxaban). N means number of animals. (Kruskal-Wallis test with Dunn’s multiple comparisons test) (The same control data were used for ONO-1600586 and rivaroxaban) **B**. PT ratio 75 minutes (before thrombus formation and bleeding test) and 4.5 hours (3 hours after thrombus formation) after oral administration of solvent (control) and FXIa (ONO-1600586) and FX inhibitors (Rivaroxaban). N means number of animals. (Kruskal-Wallis test with Dunn’s multiple comparisons test). (The same control data were used for ONO-1600586 and rivaroxaban)

### Oral administration of FXIa and FXa inhibitors suppress venous thrombus formation in rabbits

Venous thrombus formation was induced by endothelial denudation and luminal stenosis in the rabbit jugular vein after oral administration of the solvent, an FXIa inhibitor (ONO-1600586), and an FXa inhibitor (rivaroxaban). Figure 3A shows the representative macroscopic images of rabbit jugular vein thrombi in the control, FXIa inhibitor, and FXa inhibitor groups. The post-fixed thrombi were dark-reddish. Oral administration of ONO-1600586 (50 mg/kg) and rivaroxaban (15 mg/kg) significantly reduced the venous thrombus weight compared with that of the control (Figures 3A and B). Doses of 5 and 15 mg/kg ONO-1600586 and 1 and 5 mg/kg rivaroxaban tended to reduce the venous thrombus weight; however, this was not statistically significant.

**Figure 3.**
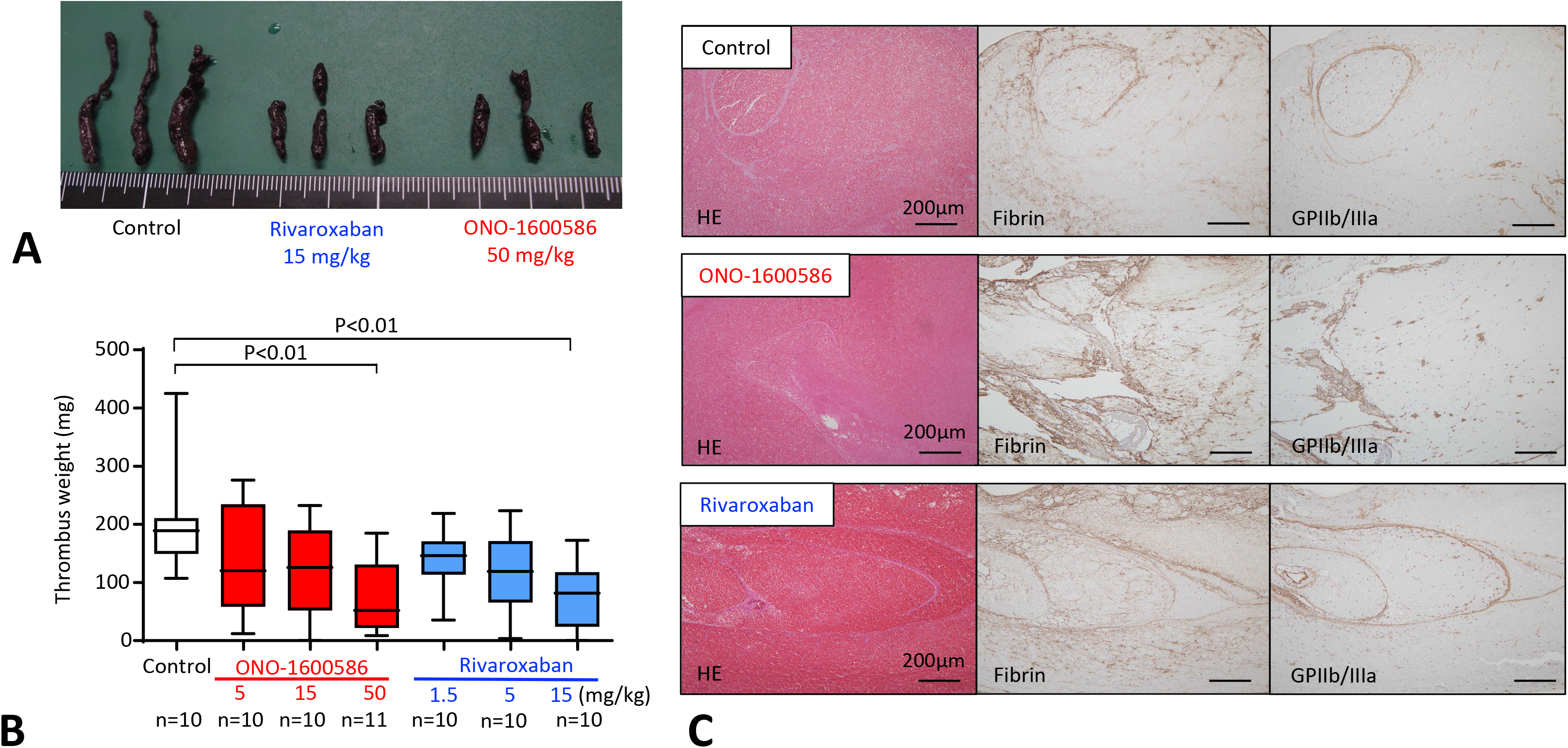
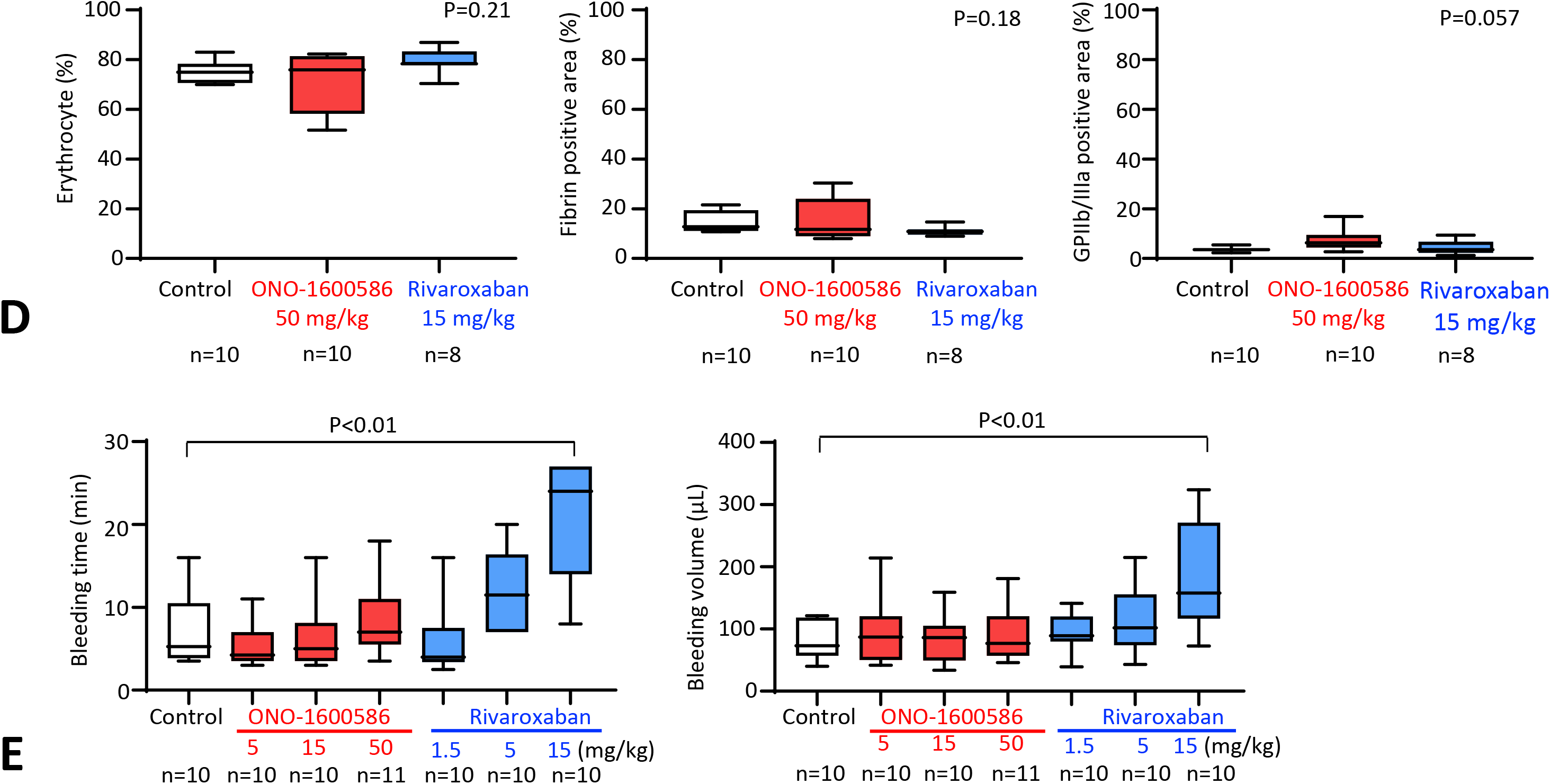
Different contribution of activated factor XI (FXIa) and activated factor X (FXa) on venous thrombus formation and skin bleeding in rabbits. **A**. Representative macroscopic images of the post-fixed venous thrombi. Thrombi are dark reddish in color in each group. Thrombi of rivaroxaban and ONO-1600586 administration groups are smaller than those of the control group. **B**. Weight of unfixed venous thrombus in control, ONO-1600586, or rivaroxaban administration groups. Oral administration of ONO-1600586 (50 mg/kg) and rivaroxaban (15 mg/kg) similarly reduced venous thrombus weight compared with that of control. Each group was compared with the control group. N means number of animals. (Kruskal-Wallis test with Dunn’s multiple comparison test) **C**. Representative histological and immunohistochemical images of venous thrombus in control, ONO-1600586, or rivaroxaban groups. All thrombi are rich in erythrocytes and contain fibrin and platelets (glycoprotein IIb/IIIa [GPIIb/IIIa]). **D**. The areas of erythrocytes or immunopositive areas for fibrin or GPIIb/IIIa in venous thrombus. N means number of histological sections. (Kruskal-Wallis test) **E**. Bleeding time and bleeding volume after oral administration of solvent (control), ONO-1600586, and rivaroxaban. Rivaroxaban dose-dependently prolonged bleeding time (left) and increased bleeding volume (right). Oral administration of ONO-1600586 did not affect bleeding time or volume. Each group was compared with the control group. N means number of animals. (Kruskal-Wallis test with Dunn’s multiple comparisons test)

Next, we histologically examined whether FXIa and FXa inhibition affected thrombus composition. All venous thrombi comprised erythrocytes, fibrin, and platelets (Figure 3C). Although the immunopositive area for fibrin tended to be smaller in the rivaroxaban group than those in the control and ONO-1600586 groups, the areas of erythrocytes or immunopositive areas for fibrin or GPIIb/IIIa did not differ among the groups (Figure 3D).

### Oral administration of FXIa inhibitor does not affect bleeding volume and time in rabbits

Skin bleeding was induced by an incision in both legs of the rabbits after oral administration of the solvent, an FXIa inhibitor, and an FXa inhibitor. However, the oral administration of ONO-1600586 did not affect the bleeding time or volume. Oral administration of rivaroxaban prolonged bleeding time and increased bleeding volume dose-dependently. Furthermore, a dose of 15 mg/kg rivaroxaban significantly prolonged the bleeding time and increased the bleeding volume compared with those of the control (Figure 3E).

### Contribution of FXIa and FXa to fibrin formation differs in the *ex vivo* blood flow chamber system

We examined the effect of FXIa and FXa on thrombus formation under low-shear conditions using a flow chamber system. Rabbit blood samples were obtained 75 min after the oral administration of the solvent, an FXIa inhibitor (ONO-1600586, 50 mg/kg), and an FXa inhibitor (rivaroxaban, 15 mg/kg). The oral administration of these doses of the inhibitors comparably inhibited venous thrombus formation in rabbits. Additionally, the surface-covering areas time-dependently increased in the three groups but did not differ among the groups (Figures 4A and B). After perfusion, we stained the thrombus on the coverslips using an anti-fibrin antibody. Immunostaining showed fibrin formation on the aggregated platelets and between the aggregated sites in the mesh pattern (Figure 4C). Furthermore, the oral administration of ONO-1600586 (50 mg/kg) significantly but incompletely inhibited fibrin formation in the flow chamber, whereas the oral administration of rivaroxaban (15 mg/kg) markedly decreased fibrin formation (Figure 4C and D). Moreover, the oral administration of ONO-1600586 and rivaroxaban did not affect neutrophil attachment on the thrombus (Figure 4E).

**Figure 4.**
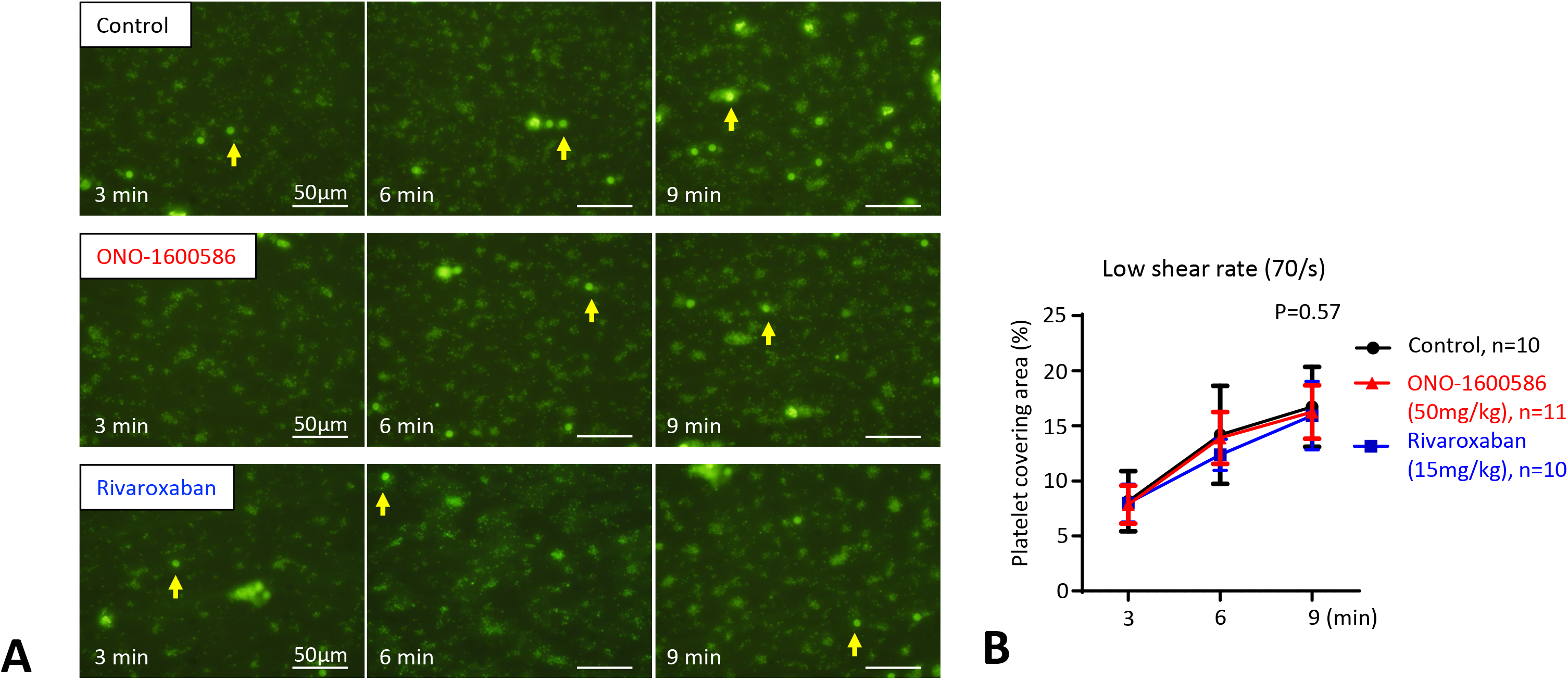
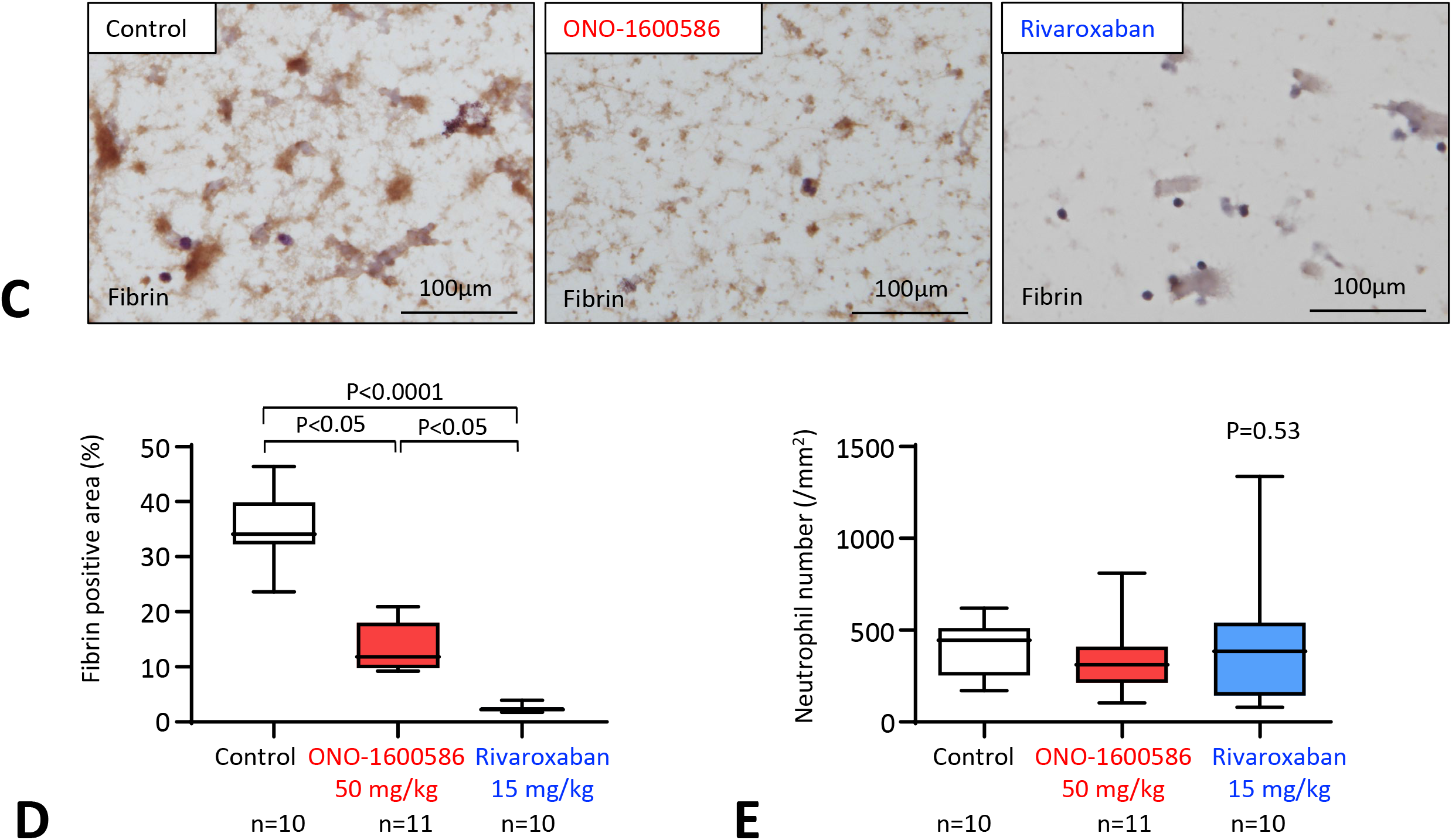
Contribution of activated factor XI (FXIa) and activated factor X (FXa) to ex vivo thrombus formation in a flow chamber system. Rabbit blood was collected 75 minutes after oral administration of solvent (control), ONO-1600586 (50 mg/kg), and rivaroxaban (15 mg/kg). The ex vivo blood was perfused in a flow chamber under low-shear rate of 70/s. **A**. Representative fluorescent images under low shear rate (70/s). Platelets and leukocytes were labeled with mepacrine, and the fluorescent images were captured 3, 6, and 9 minutes after perfusion. The surface covering area time-dependently increases in all three groups. Large fluorescent dots indicate leukocytes (arrows). **B**. The surface covering areas under low-shear rate in three groups. (Two-way repeated measure ANOVA with Tukey’s multiple comparison test) **C**. Representative immuno-staining images for fibrin on glass coverslip after perfusion. Mesh-like pattern of fibrin formation on islands of aggregated platelets and between the islands in control. Reduced fibrin formation in ONO-1600586 administration. Absence or little fibrin formation in rivaroxaban administration. **D**. Fibrin-immunopositive area on glass coverslip after ex vivo blood perfusion under low-shear rate. (Kruskal-Wallis test with Dunn’s multiple comparison test) **E**. Number of leukocyte adhesions after ex vivo blood perfusion under low-shear rate (Kruskal-Wallis test)

### ONO-1600586 suppresses human FXIa activity and reduces thrombin generation and fibrin formation *in vitro* human blood perfusion

To examine the effects of ONO-1600586 on human FXIa and thrombus formation in vitro, we measured FX and FXI activities, thrombin generation, and fibrin formation before and after human blood perfusion in a flow chamber system. The administration of the FXIa inhibitor ONO-1600586 inhibited fibrin formation but did not affect the platelet covering area and neutrophil attachment on the *in vitro* thrombus as rabbit blood perfusion in the flow chamber system (Figure 5A-D). FXIa activity, rather than FXa activity, in the plasma after perfusion was higher than that in the plasma before perfusion. However, the administration of FXIa inhibitor suppressed the enhanced FXIa activity and plasma levels of prothrombin fragment 1+2 (Figure 5E).

**Figure 5.**
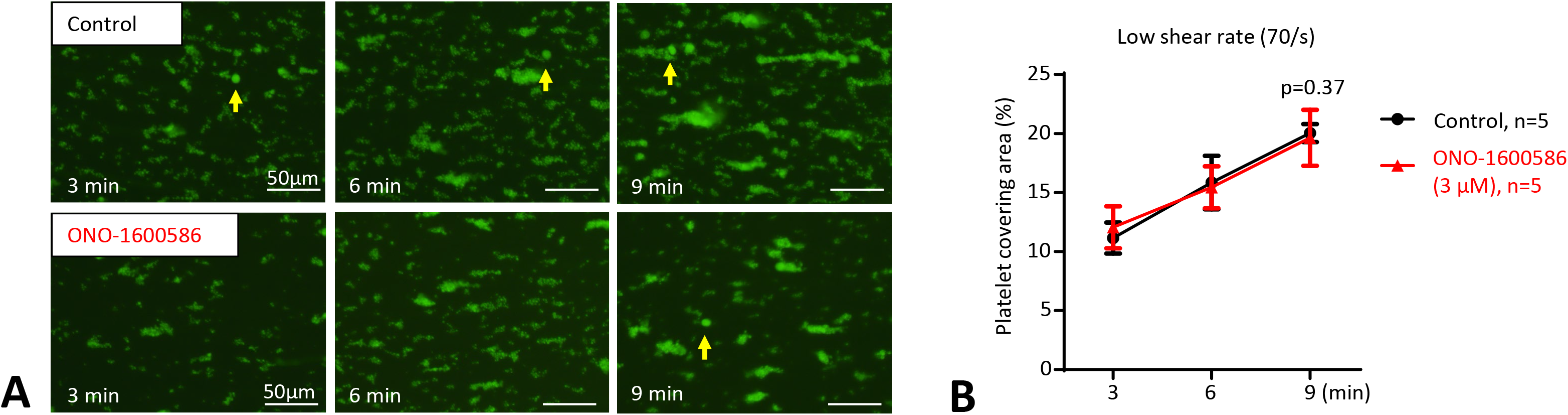
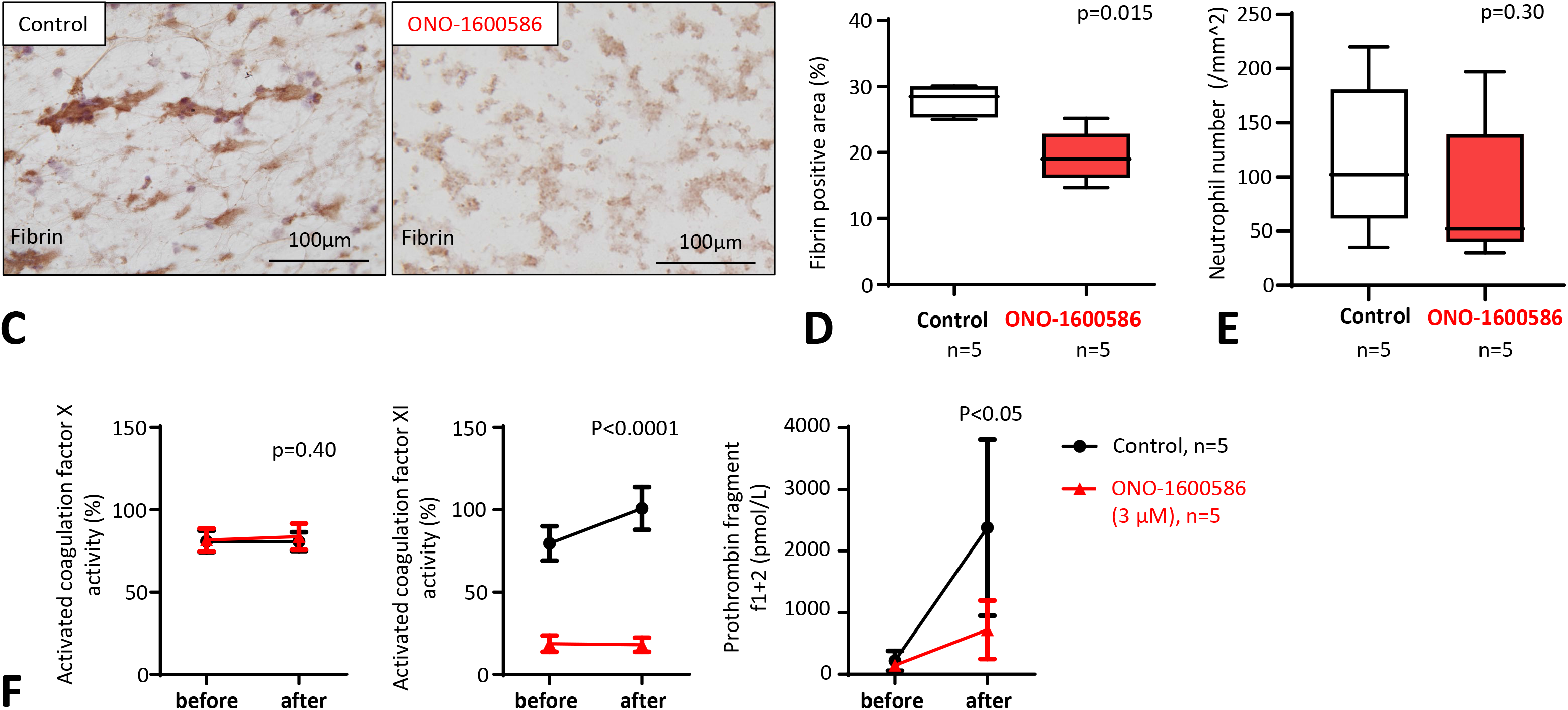
Effects of ONO-1600586 on human activated factor XI (FXIa) and in vitro thrombus formation. The human blood was perfused in a flow chamber under low-shear rate of 70/s. **A**. Representative fluorescent images under low shear rate (70/s). Platelets and leukocytes were labeled with mepacrine, and the fluorescent images were captured 3, 6, and 9 minutes after perfusion. The surface covering area time-dependently increases in all three groups. Large fluorescent dots indicate leukocytes (arrows). **B**. The surface covering areas under low-shear rate in control (solvent) and ONO-1600586 addition (Two-way repeated measure ANOVA with Tukey’s multiple comparisons test) **C**. Representative immuno-staining images for fibrin on the glass coverslips after perfusion. ONO-1600586 addition decreases fibrin formation. **D**. Fibrin-immunopositive area after human blood perfusion under low-shear rate (Mann-Whitney U-test) **E**. Number of leukocyte adhesion after human blood perfusion under low-shear rate (Mann-Whitney U-test) **F**. Activated factor X (FXa) activity, FXIa activity, levels of prothrombin fragment f1+2 before and after perfusion of human blood. (Two-way repeated measure ANOVA with Tukey’s multiple comparison test)

## Discussion

This study showed that FXI was mainly localized in the non-organizing area of human DVT and was closely distributed in fibrin. Additionally, an orally administered direct FXIa inhibitor, ONO-1600586, rather than the FXa inhibitor, did not enhance bleeding under suppressed venous thrombus formation in rabbits, whereas ONO-1600586 incompletely inhibited fibrin formation *ex vivo* under low-shear conditions. ONO-1600586 suppressed human FXIa activity and reduced thrombin generation and fibrin formation during human blood perfusion.

FXI was present in all aspirated DVT, mainly localized in fibrin– and erythrocyte-rich non-organizing areas and closely distributed in fibrin. The presence of intrinsic coagulation factors in human DVT has been reported previously. Sugita et al. examined the localization of factor VIII (FVIII), VWF, and fibrin in DVT and found co-localization of FVIII with platelets, VWF, and fibrin [17]. The results suggest that the localization of FXI in DVT is not always similar to that of FVIII. FXI circulates in the plasma, is stored in platelets, and is secreted upon platelet activation [18]. Although FXIa activity and antigen have been described in washed platelet suspensions, it has been estimated that FXIa activity constitutes approximately 0.5% of the FXIa activity in normal plasma [18]. Additionally, the plasma level of FXI is associated with DVT incidence [4–7]. Predominant co-localization with fibrin is comparable to these studies, and FXI was abundant in the non-organizing areas compared with the organizing area in this study. Our findings suggest that FXI participates in human DVT formation and may be degraded during the organizing process, similar to other thrombus components, such as erythrocytes, fibrin, platelets, and neutrophils.

The direct oral FXIa inhibitor ONO-1600586 suppressed endothelial denudation and stenosis-induced venous thrombus formation without excessive bleeding in rabbits. Notably, these findings are comparable with those in FXI-deficient mice [8], anti-FXI antibodies [11,19,20], FXI antisense oligonucleotides [10,21], and small molecule FXIa inhibitors [9]. However, ferric chloride [8,10] and silk thread [9] are not physiological initiators of venous thrombus formation; therefore, the data should be interpreted cautiously. Our study strongly supports the hypothesis that the enzymatic activity of FXIa contributes to venous thrombus formation *in vivo*. The relative areas of erythrocytes, fibrin, and platelets in the venous thrombi did not differ between groups, suggesting that the compositions are independent of the venous thrombus size *in vivo*. A previous study showed that an anti-FXI antibody reduced both platelet and fibrin contents in venous thrombi compared with controls [11]. The difference might be because of FXIa inhibition rather than FXI inhibition.

Orally available small-molecule inhibitors are more suitable for the long-term prevention of venous thrombosis than large-molecule inhibitors. Intravenous administration of BMS-262084, which is an irreversible small-molecule FXIa inhibitor, reduced silk thread-induced thrombus in the inferior vena cava of rabbits [9]. Therefore, this study is the first to show that oral administration of an FXIa inhibitor is effective for venous thrombus formation *in vivo*. Although ONO-1600586 can inhibit plasma kallikrein, the higher IC50 value of human kallikrein compared with that of human FXIa suggests that the antithrombotic effect mostly reflects those of FXIa inhibition in rabbits.

Clinical and experimental studies have shown that FXI deficiency and inhibition have a lower bleeding tendency than FX deficiency and inhibition [6,7,22]. In fact, FXIa and FXa inhibition showed different hemostatic effects with similarly reduced venous thrombus formation in this study. These results are consistent with those of a previous study [20]. Anti-human FXI antibodies mildly inhibited ferric chloride-induced venous thrombus formation without altering tail-cut bleeding time in mice, whereas enoxaparin strongly inhibited venous thrombus formation and prolonged bleeding time [20]. However, why FXI/FXIa inhibition does not result in excess bleeding remains unclear. Our *ex vivo* study showed that FXIa inhibition significantly but incompletely inhibited mural fibrin formation under low-shear conditions compared with FXa inhibition (Figures 4C and D). Therefore, the limited contribution of fibrin formation on the vascular surface may explain the differences in bleeding tendencies between FXI and FX deficiency and inhibition.

Oral and parenteral FXI/FXa inhibitors have been evaluated in phase I and II clinical trials and are undergoing phase III evaluation. The present FXIa inhibitor ONO-1600586 suppressed human FXIa activity, thrombin generation, and fibrin formation under low-shear conditions (Figure 5A). These results suggest that human FXI can be activated via thrombin during venous thrombus formation and that the small molecular inhibitor ONO-1600586 can inhibit FXIa activity under low-shear conditions. Phase I study on ONO-1600586-related product, ONO-7684, which is an oral FXIa inhibitor, showed safety and tolerability for approximately 14 days in the fed state in healthy individuals [23]. An open-labeled non-inferiority phase II trial compared osocimab, human monoclonal anti-FXI antibody, enoxaparin, and apixaban for thromboprophylaxis in patients with who have undergone knee arthroplasty [24]. Single infusion of osocimab showed non-inferiority compared with enoxaparin for incidence of VTE at 10–13 days. The major or clinically relevant non-major bleeding rate was approximately 4.7%, 5.9%, and 2% in patients receiving osocimab, enoxaparin, and apixaban, respectively. PACIFIC-AF (Safety of the Oral Factor XIa Inhibitor Asundexian Compared With Apixaban in Patients With Atrial Fibrillation) trial compared the efficacy and safety of asundexian, which is a small molecule FXIa inhibitor, and apixaban. Although underpowered for efficacy, asundexian (50 mg, one daily) showed a lower bleeding rate (0.4%) than apixaban (2.4%) [25]. Therefore, further studies should establish the efficacy and safety of FXI/ FXIa inhibitors in humans.

This study had some limitations. First, the sample size of human-aspirated DVT was small. However, the consistent presence of FXI in fibrin-rich areas suggests that FXI plays a role in fibrin formation during DVT. CD34 is not a specific maker of endothelium. Because platelets express CD31, an endothelial marker, we used anti-CD34 antibody for endothelialization. Second, rabbit FXI lacks Cys321 and therefore is circulating as a noncovalently associated dimer [26]. This may affect functional difference between rabbit and human FXIa. However, ONO-1600586 similarly prolonged aPTT in human and rabbits.

In conclusion, FXI localization may provide a rationale for FXI/FXIa inhibition to prevent human DVT, and preserved fibrin formation under FXIa inhibition may be associated with a minor hemostatic role of FXIa.

## Acknowledgments

We thank Nahoko Udatsu and Kyoko Ohashi for their technical assistance. We would like to thank *Editage* (www.editage.com) for English language editing.

## Supplemental Material

Tables S1–S2

Figures S1–S3

## Figure Captions

**Figure S1.**
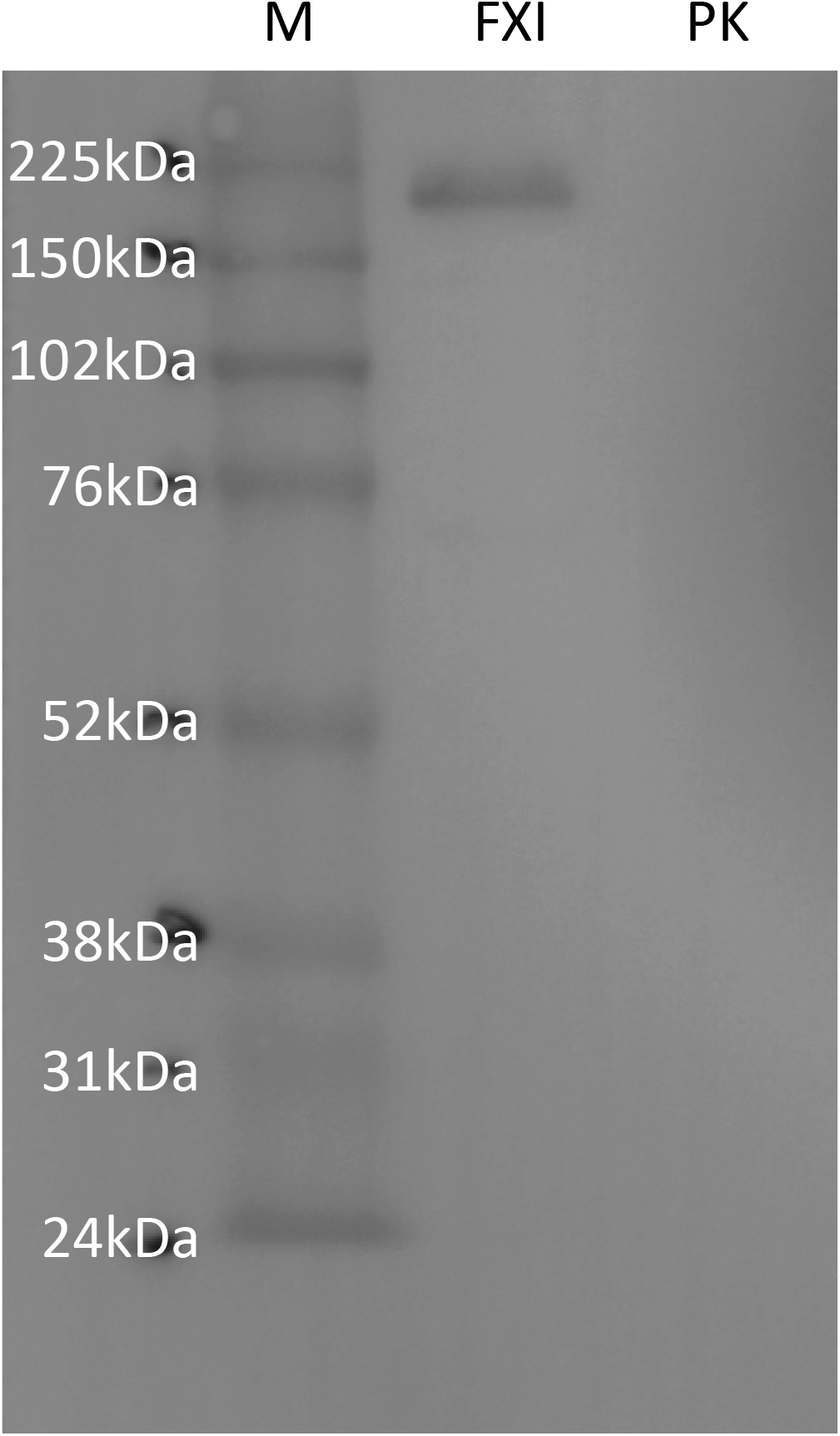
Western blot using anti-FXI antibody. M: ECL Rainbow Marker-Full range (Cytiva, Tokyo, Japan) FXI: purified human FXI (HFXI 1111, Enzyme Research Laboratories Ltd., Swansea, UK); PK: purified human prekallikrein (HPK 1302, Enzyme Research Laboratories Ltd.) The western blotting using anti-sheep FXI antibody (sheep polyclonal, LS-B10243; LifeSpan, Inc. Seattle, USA, final concentration 10µg/mL) was performed under non-reducing condition.

**Figure S2.**
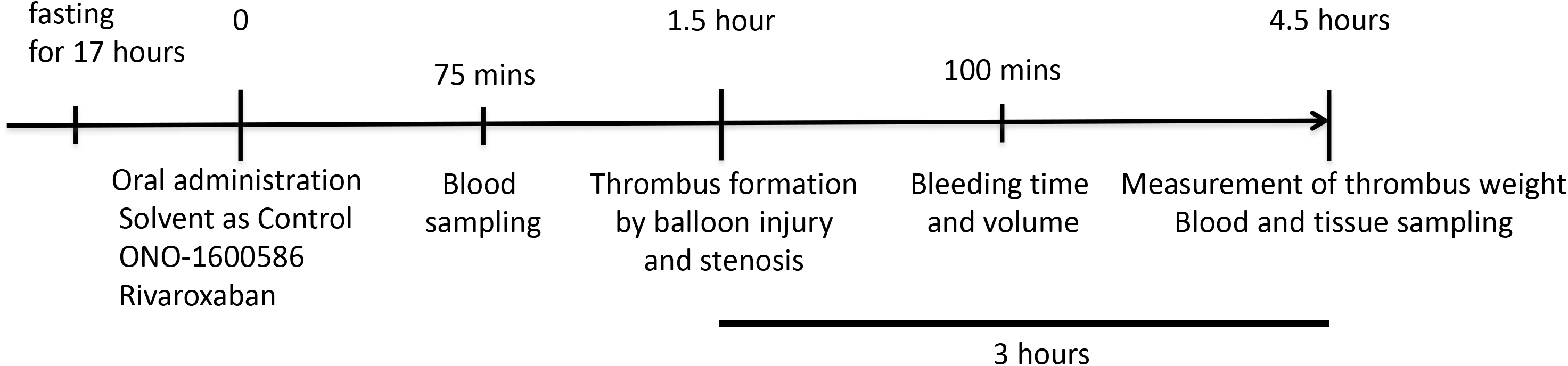
A protocol of rabbit model of jugular vein thrombosis and skin bleeding. ONO-1600586, an activated factor XI inhibitor

**Figure S3.**
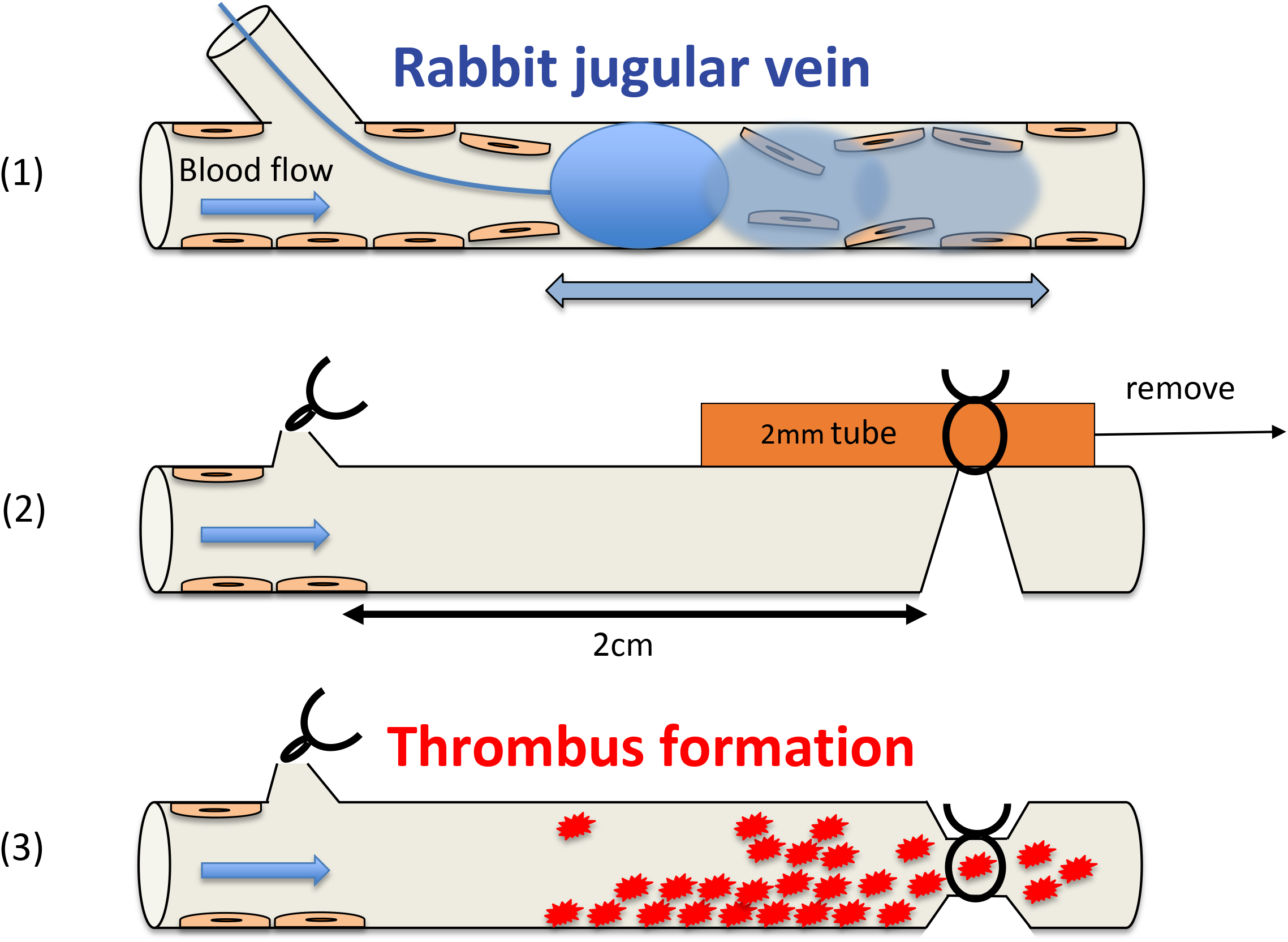
Procedure of rabbit jugular vein thrombus model. (1) Balloon catheter (3F) inserted 5 cm into jugular vein via distal branch was inflated three times (airvolume,0.4 mL) for endothelial denudation. (2) For induce luminal stenosis at jugular vein 2cm-proximal from the branch, both jugular vein and a polyethylene tube of 2 mm in outer diameter were ligated, and the tube was removed. (3) Thrombi were sampled and weighed 3 hours after thrombus formation (4.5 hours after administration of solvent or inhibitors).

**Table S1.** Clinical background of DVT patients (n = 15)

|  |  |  |
| --- | --- | --- |
| Age, median (range; years) | 54 | (20-78) |
| Male sex, n (%) | 8 | (53%) |
| Obesity; BMI >25, n (%) | 3 | (20%) |
| Smoking, n (%) | 8 | (53%) |
| Complication |  |  |
| Posttraumatic, n (%) | 4 | (27%) |
| Cancer, n (%) | 2 | (13%) |
| Antiphospholipid syndrome, n (%) | 0 | (0%) |
| Medication |  |  |
| Aspirin, n (%) | 2 | (13%) |
| Warfarin, n (%) | 3 | (20%) |
| Chemotherapy, n (%) | 0 | (0%) |
| Steroid, n (%) | 3 | (20%) |
| Treatment period of heparin infusion<br>before aspiration, median (range; days) | 4 | (0-30) |
| Laboratory data before thrombus aspiration, median (range) |  |  |
| Complete blood cell count |  |  |
| White blood cell (x10<sup>3</sup>/μL) | 6.8 | (4.1-14.4) |
| Red blood cell (x10<sup>6</sup>/μL) | 3.68 | (2.3-4.88) |
| Hemoglobin (g/dL) | 11.8 | (7.5-17.1) |
| Platelet (x10<sup>3</sup>/μL) | 225 | (117-724) |
| Coagulation |  |  |
| D-dimer (μg/mL) | 13 | (4.5-76.2) |
| PT-INR (INR) | 1.11 | (0.9-2.1) |
| aPTT (sec.) | 31.7 | (23.5-45.2) |
| Activated protein S (%) | 78 | (28-144) |
| Activated protein C (%) | 96 | (29-134) |
| Antithrombin III activity (%; n=14) | 91 | (35-126) |

aPTT, activated partial thromboplastin time; DVT, deep vein thrombosis;
PT-INR, prothrombin time-international normalized ratio;

**Table S2.** Effects of ONO-1600586 on human coagulation enzymes.

| Human enzymes | IC50 (μM) |
| --- | --- |
| Activated factor XI | 0.0020 |
| Plasma Kallikrein | 0.12 |
| Thrombin | 26 |
| Activated factor VII | >33 |
| Activated factor IX | >100 |
| Activated factor XII | >33 |
| Plasmin | 67 |
| Urokinase | >33 |
| Tissue plasminogen activator | >33 |

